# Enabling the Study of Gene Function in Gymnosperms: VIGS in *Ephedra tweedieana*

**DOI:** 10.1101/2025.02.07.637168

**Authors:** Anthony Garcia, Jo Trang Bùi, Todd P. Michael, Stefanie M. Ickert-Bond, Verónica S. Di Stilio

**Affiliations:** University of Washington, Department of Biology, Seattle, WA, USA; Salk Institute, La Jolla, CA, USA; University of Alaska, Fairbanks, AK, USA

**Keywords:** gymnosperm, Gnetales, VIGS, PDS, functional studies, gene silencing, transformation, TRV

## Abstract

**Premise:** As the sister clade to angiosperms, gymnosperms are key to enabling the reconstruction of ancestral gene regulatory networks for seed plants. However, tools to rapidly and efficiently investigate gene function in gymnosperms remain limited due to the challenges of long life cycles and large genome sizes. Species within the xerophytic genus *Ephedra* (Gnetales) have comparatively smaller genomes and shrubby growth habits with shorter life spans, making them better suited for greenhouse cultivation and laboratory experiments.

**Methods and Results:** Here, we implement Virus-Induced Gene Silencing (VIGS) to manipulate gene expression in *Ephedra tweedieana. Agrobacterium*-mediated vacuum infiltration of Tobacco Rattle Virus (TRV2 and TRV1) in seedlings resulted in highly efficient silencing of the *E. tweedieana PHYTOENE DESATURASE* ortholog *EtwPDS*. The expected photobleaching phenotype was observed as early as two weeks. It lasted at least three months, in stems, shoot tips, leaves, axillary meristems, and lateral branches of treated plants.

**Conclusions:** This first report of transient transformation and targeted gene silencing in a gymnosperm will further enable functional studies of the genetic mechanisms underpinning adaptations in this important and underrepresented lineage of seed plants.

## INTRODUCTION

Gymnosperms abound in present-day ecosystems, including some of the most important timber species (e.g., pine, spruce, and fir), and the largest (giant redwoods) and oldest (bristlecone pine) species on earth. *Ephedra* L. comprises approximately 54 species distributed worldwide, New World species occur in the North American deserts of the southwest and arid and semi-arid regions of South America (Ickert-Bond and Renner, 2016). Many aspects of diversification within *Ephedra* remain unresolved as a result of widespread polyploidy and the paucity of comparative data. Along with two other disparate genera, *Gnetum* L. and *Welwitschia* Hook.f., they form the gymnosperm order Gnetales nested within the conifers, which together with cycads, ginkgoes, and flowering plants (angiosperms) comprises the seed plants. Gnetales have interesting convergences with the flowering plants (angiosperms), including double fertilization and flower-like reproductive structures (Bowe et al., 2000), and the occasional occurrence of structurally bisexual cones (Ickert-Bond and Renner, 2016).

Climate is shifting towards warmer temperatures and more weather extremes, threatening agriculture worldwide (Heino et al., 2023). *Ephedra*’s success has been largely attributed to its ability to remain metabolically active year-round, with photosynthetic stems, highly reduced ephemeral leaves, sunken stomata, and large taproots, becoming the dominant element of the flora in extreme environments (high altitude and arid ecosystems; Ickert-Bond and Renner, 2016). Research on the genetic and developmental basis of adaptations to extreme environments and of seed plant character evolution has been hindered by a lack of gymnosperm model systems since most are trees with decades-long generation times. Having relatively small genomes, 7-8 giga base pairs (Gbp), and shorter generation times (as short as 2 years), *Ephedra* is amenable to investigations on the evolution of gymnosperm and seed plant innovations. Uncovering the genes underlying key innovations in this lineage requires a genetic toolkit including transformation protocols for functional studies.

Virus-Induced Gene Silencing (VIGS) has proven successful as a transient transformation technique for evo-devo studies across a variety of plants (Di Stilio, 2011), but still remains limited to angiosperms (Lange et al., 2013; Dommes et al., 2019; Rössner et al., 2022). In bryophytes, stable transformation techniques are facilitated by their haploid, gametophyte-dominant life cycles (Yadav et al., 2023). In the model fern *Ceratopteris richardii* Brongn., particle bombardment (Rutherford et al., 2004; Plackett et al., 2014) and *Agrobacterium*-mediated transformation of spores (Muthukumar et al., 2013) and gametophytes (Jiang et al., 2024) have facilitated functional gene studies via stable transformation. In gymnosperms, transformation has been achieved in conifers and *Ginkgo biloba* L. (ginkgo). Stable and transient transformation in conifers relies on particle bombardment or *Agrobacterium*-mediated transformation and regeneration (Morris et al., 1989; Moyle et al., 2002; Wagner et al., 2005; Tahir et al., 2011; Perez-Matas et al., 2023; Zhao et al., 2024). In ginkgo, *Agrobacterium-* mediated stable transformation (Dupré et al., 2000; Ayadi and Trémouillaux-Guiller, 2003) and protoplast transient transformation (Han et al., 2023) have also been described. However, the challenges of long life cycles and complicated protocols have limited the transformation of those species to forestry and industry applications. VIGS, on the other hand, has not been previously described in a gymnosperm and would allow for faster and more cost-effective functional analyses. Here, we describe a method for efficient virus-mediated transient transformation in *Ephedra tweedieana* Fisch. & C.A.Mey., a key tool in developing a gymnosperm model organism.

## METHODS

### Plant cultivation

*Ephedra tweedieana* seeds (from a population near Montevideo, Uruguay) were germinated on 1% Agar plates in growth chambers under 50% humidity, 120 µm light (Red: Far-Red ratio =1.0), and 16 hr. light/8 hr. dark cycle. Germination success was 93% (51/55 seeds). After the emergence of a taproot, seedlings were transplanted to soil (Sunshine Mix #4, Sun Gro Horticulture, Agawam, MA) and kept in growth chambers under the above conditions, with 85% survival rate (47 seedlings).

### *Identification of the* Ephedra tweedieana PHYTOENE DESATURASE *ortholog* EtwPDS

To identify the *PHYTOENE DESATURASE* ortholog, we queried the *E. tweedieana* proteome with the *Arabidopsis PHYTOENE DESATURASE* (AT4G14210) PDS3 coding sequence. The proteome was first searched using a gene family analysis (Orthofinder) based on closely related gymnosperm and model plant proteomes (resources.michael.salk.edu/resources/ephedra_genomes/). Then, the *E. tweedieana* proteome was searched using BLASTP to ensure that all potential orthologs were identified. Two potential orthologs were identified on Chromosome 7 (Chr07) and Chr03. Multiple sequencing alignment (MSA) revealed that the ortholog on Chr07 was truncated. Therefore, we utilized the ortholog on Chr03: EtweTM011.hap2.chr3.a04.g228080.t1 to build our construct (Fig. S1) and named it Etw*PDS*.

### Generation of Tobacco Rattle Virus (TRV) constructs

A 425 base-pair fragment of the *EtwPDS* gene was amplified from cDNA with locus-specific primers (Table S1), digested with KpnI-HF and XhoI (New England Biolabs, Ipswich, MA, USA) and ligated into pYL156 (TRV2, Addgene plasmid #148969, Liu et al., 2002) using T4 DNA Ligase (New England Biolabs, Ipswich, MA, USA). Three clones were verified by Sanger sequencing (Genewiz, South Plainfield, NJ, USA). *Agrobacterium tumefaciens* GV3101 was transformed by electroporation with TRV2-*EtwPDS*, empty TRV2 (EV) or TRV1 (pYL192, Addgene plasmid # 148968, Liu et al., 2002), and the resulting colonies were confirmed by PCR.

### Agroinfiltration of Ephedra tweedieana seedlings

Virus-induced gene silencing of *EtwPDS* was performed on one to two-week-old seedlings as previously described for *Thalictrum dioicum* L. (Di Stilio et al., 2010) with the following modifications: 10 mL of 0.5 M MES (2-(4-Morpholino)-Ethane Sulfonic Acid) was used instead of 5 mL 1.0 M MES and two vacuum infiltration regimes of 2 and 5 min instead of a single infiltration time of 2 minutes. In addition to VIGS-treated plants (n=21), we included two negative controls: TRV1 + TRV2 empty vector (EV, n=10) to control for virus infection effects, and infiltration media (IM, n=3) to control for the vacuum infiltration treatment. Treated seedlings were returned to pots containing Sunshine#4 soil, covered with a plastic dome, and allowed to recover and grow under the conditions specified above.

### Validation of TRV infection and gene silencing

Three weeks following VIGS treatment, approximately 10 mg of cotyledon or stem tissue was collected and flash frozen in liquid nitrogen, and RNA was extracted from using Direct-zol RNA Miniprep Kit (Zymo Research, Irvine, CA, USA). First-strand synthesis was carried out with iScript™ cDNA Synthesis Kit (Bio-Rad, Laboratories, Hercules, CA, USA) from 1 µg of RNA and used in reverse transcription PCR (RT-PCR) for TRV RNA detection and in real-time quantitative PCR (qPCR) for the validation of gene expression.

RT-PCR was conducted to detect TRV1, TRV2, and the EF1b reference gene with locus-specific primers (Table S1). Samples were amplified in a thermocycler (DNA Engine DYAD Peltier Thermal Cycler, MJ Research, St. Bruno, QC, Canada) using the following conditions: initial denaturation 2 min at 95°C, 30 cycles of denaturation for 30 sec at 95°C, annealing for 30 sec at 51°C, and extension for 1 min at 72°C, followed by a final extension at 72°C for 5 min. Samples were loaded on a 1% TAE gel, run at 100V for 60 minutes, and visualized using the Bio-Rad Gel Doc EZ Imager (Bio-Rad Laboratories, Hercules, CA, USA).

qPCR was conducted using the iTaq Universal SYBR Green Supermix and gene-specific primers to detect target *EtwPDS* and housekeeping *EtwEF1b* expression levels (Supplemental Table 1). Samples were run using a Bio-Rad CFX Connect Real-Time System (Bio-Rad Laboratories, Hercules, CA, USA) set to the following conditions: initial denaturation for 5 min at 94°C, 40 cycles of denaturation for 30 sec at 94°C, annealing for 30 sec at 57°C, extension for 30 sec at 72°C, and plate read, with cycling followed by melt curve analysis increasing in 0.5°C every 5 sec from 65°C to 95°C. *EtwPDS* expression was normalized to *EtwEF1b* using the delta delta CT method (Livak and Schmittgen, 2001), and a one-way ANOVA was performed to compare expression levels of treatments and controls.

### Imaging

Plants were photographed using a Nikon D3400 hand-held digital camera or a dissecting microscope (Nikon SMZ800, Nikon Instruments Inc., Melville, NY, USA) equipped with a QImaging MicroPublisher 3.3 RTV digital camera (Surrey, BC, Canada). Images were edited and compiled using Affinity Publisher v1.10.6 and Inkscape v1.4.

## RESULTS

### Vacuum Agrobacterium infiltration triggers TRV infection and targeted gene silencing

To enable the investigation of gene function in the gymnosperm *E. tweedieana*, we aimed to obtain proof-of-principle evidence of transient transformation by Virus-Induced Gene Silencing (VIGS). To that end, we tested whether *Agrobacterium*-mediated infiltration of Tobacco Rattle Virus (TRV) can trigger targeted gene silencing of the commonly used marker *PHYTOENE DESATURASE* resulting in photobleaching of green photosynthetic tissues (Figure 1). Vacuum infiltration of seedlings with *Agrobacterium tumefaciens* carrying *E. tweedieana PHYTOENE DESATURASE (*TRV2-*EtwPDS)* resulted in high rates of viral infection, between 30% and 80% depending on the specific treatment (Table 1).

**Figure 1:**
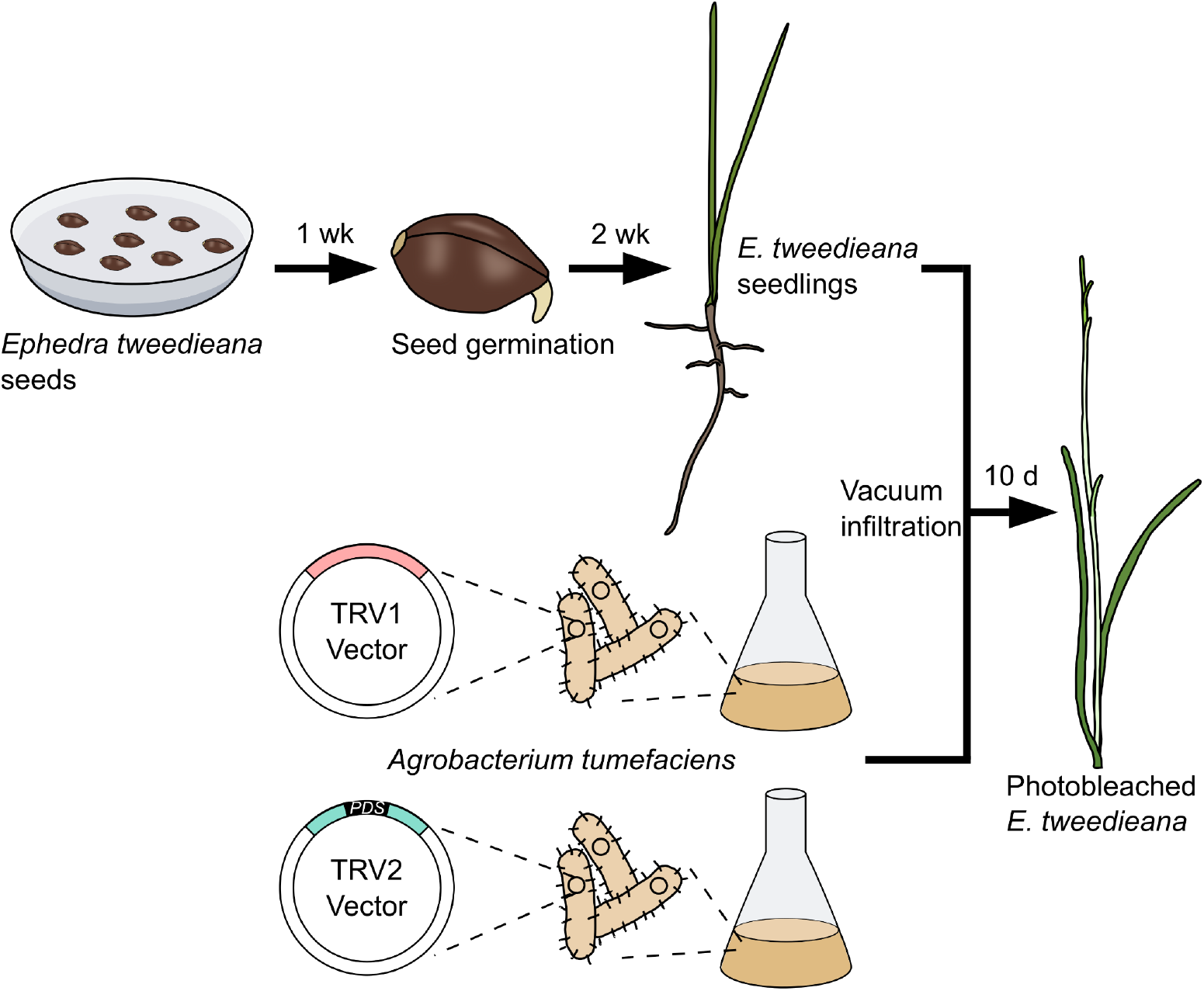
VIGS workflow in the gymnosperm *Ephedra tweedieana*. *Agrobacterium-*mediated infiltration of TRV in *E. tweedieana* seedlings triggers virus-induced gene silencing of the *PHYTOENE DESATURASE* ortholog Etw*PDS*. Seeds germinate on agar after one week and are transplanted to soil. Two-week old seedlings are infiltrated under vacuum for two minutes with equal parts of *Agrobacteria* cultures carrying TRV1 and TRV2 vectors, the latter with a fragment of the target gene. Photobleaching of green tissues, a sign of Etw*PDS* silencing, was observed ten days after treatment.

**Table 1:** Summary statistics for VIGS of the *PHYTOENE DESATURASE* ortholog in *Ephedra tweedieana*. Molecular validation via the detection of Tobacco Rattle Virus (TRV) RNA and photobleaching phenotype in different tissues of controls (infiltration media and empty vector, EV) and of plants treated under two or five-min vacuum infiltration.

| Treatment | Plants treated | Validation |  | Photobleaching |  |  |  |
| --- | --- | --- | --- | --- | --- | --- | --- |
|  |  | TRV (%) | silenced (%) | Internode (%) | Lateral shoot (%) | Apical node (%) | Leaf (%) |
| Infiltration Media Control | 3 | 0 | NA | 0 | 0 | 0 | 0 |
| TRV2-EV | 10 | 8 (80%) | 0 | 0 | 0 | 0 | 0 |
| TRV2-EtwPDS (2 min) | 11 | 7 (64%) | 7 (64%) | 7 (64%) | 2 (18%) | 2 (18%) | 6 (55%) |
| TRV2-EtwPDS (5 min) | 10 | 4 (40%) | 3 (30%) | 3 (30%) | 0 | 2 (20%) | 2 (20%) |

Infection by TRV, validated by amplification of viral RNA, was only observed in plants treated with TRV1 and TRV2, not in infiltration media (IM) control plants (Fig. 2g). Plants treated with an empty vector, TRV2-EV, were no different than IM control plants in leaf morphology, shoot morphology, or growth habit, suggesting *Agrobacterium* infection with TRV alone does not alter *E. tweedieana* growth and development (Supplemental Figure 1). Vacuum infiltration for two minutes resulted in a higher infection rate (64%) than five minutes (40%, Table 1), hence longer infiltration times are not warranted, nor beneficial for increasing infiltration rate.

**Figure 2:**
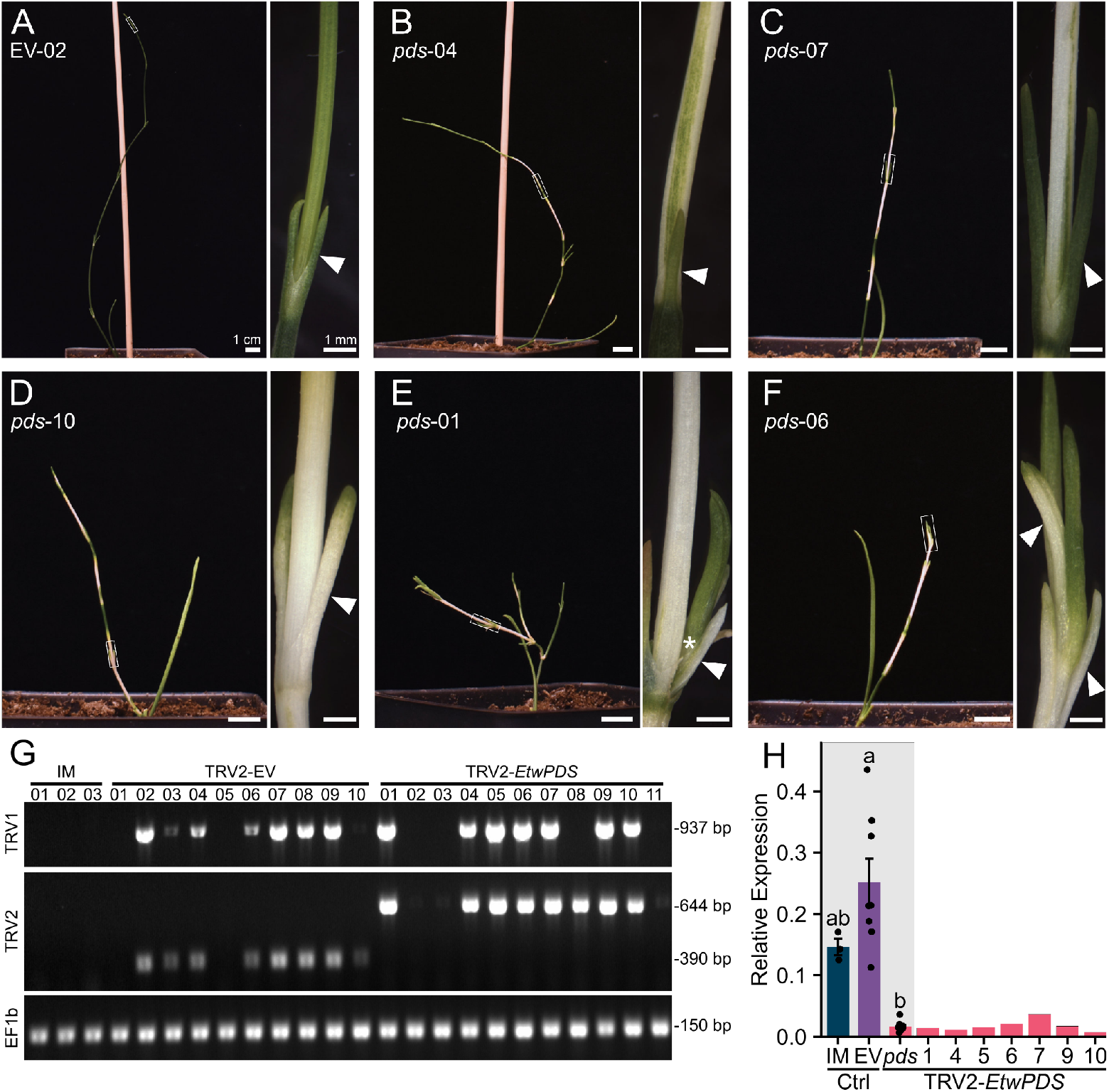
Photobleaching in *Ephedra tweedieana* by Virus-induced Gene Silencing. Full plant images show a range of bleaching severity, and the corresponding narrow panel displays magnified detail of the region marked in a white box. a) Empty-vector control plant. b-f) Representative TRV*2*-*EtwPDS* treated plants displaying a range of phenotypes. Scale bars 1 cm, or 1 mm (insets). Arrowheads point to leaves, and asterisk indicates lateral branch. g) RT-PCR validation of TRV1 RNA and TRV2 RNA presence, including the reference gene *EF1b* as a loading control. h) qPCR validation of *PDS* silencing in bleached tissues of virus-validated plants, relative to EF1b. Different letters indicate statistical significance (*P*<0.05) in a one-way ANOVA (*F*2,15 = 18.42, *P* = 9.13e-05) followed by a Tukey HSD post-hoc test (*P* _EV vs IM_ = 0.126, *P*_EV vs *pds*_ = 0.00006, *P*_IM vs *pds*_ = 0.06).

Plants treated with TRV2*-EtwPDS* experienced down-regulation of *EtwPDS*, to an approximately 10-fold decrease in *EtwPDS* expression compared to TRV2-EV controls (Figure 2h). One 5-minute treated plant showed similar levels of *EtwPDS* expression to TRV-EV plants (Supplemental Figure 2). Taken together, our results indicate that *Agrobacterium-*mediated infiltration of TRV causes TRV infection that triggers gene silencing in *E. tweedieana*.

### Ephedra tweedieana PDS *silencing causes photobleaching in a range of tissues*

*PDS* silencing typically results in photobleaching of photosynthetic tissue by disrupting carotenoid biosynthesis (Wang et al., 2009) and is commonly used as a visual marker in targeted gene silencing experiments. Photobleaching of *E. tweedieana* tissues was first observed 10 d after infiltration in TRV-*EtwPDS* silenced plants (Table 1, Supplemental Figure 3). Three weeks following treatment, a range of photobleaching was observed compared to controls, with some plants overcoming silencing (Figure 1a-c) while others displayed stunted growth and severe silencing (Figure 2f).

To determine the range of tissues susceptible to VIGS in *E. tweedieana*, we examined specific tissues affected by photobleaching by light microscopy (Table 1). All plants undergoing silencing of *EtwPDS* exhibited photobleaching of stem internodes (Figure 2f), and most also showed leaf photobleaching. Additionally, two plants showed photobleaching of lateral shoots (Figure 2e) and apical leaves (Figure 2f). This suggests that, while stem internodes were most susceptible to VIGS, gene silencing can also encompass lateral and apical meristematic tissue, extending the phenotypic effects to tissues that develop after treatment.

## DISCUSSION

To the best of our knowledge, this is the first report of VIGS in a non-flowering plant. The gene silencing efficiency of 64 % we observed in *E. tweedieana* is comparable to or higher than in other species where VIGS has been implemented. This includes several species in the Ranunculales: *Eschscholzia californica* Cham. (up to 94%, Wege et al., 2007), *Thalictrum dioicum* (42%, Di Stilio et al., 2010), *Papaver somniferum* L. (23%, Hileman et al., 2005), and *Aquilegia coerulea* E.James (up to 12%, Gould and Kramer, 2007; Sharma and Kramer, 2013) and across eudicots and monocots (Di Stilio, 2011).

As the first described example of VIGS in a gymnosperm species, the rapid and high efficiency of silencing by VIGS in *Ephedra* will facilitate testing of candidate genes to investigate the evolution of gene function, bridging the gap across seed plants and land plants more broadly. This tool also opens the door for studying the wealth of gymnosperm diversity across clades such as cycads and Gnetales. Within *Ephedra*, a number of avenues exist for studying stress response and convergent evolution of fleshy reproductive structures analogous to angiosperm fruits (Di Stilio and Ickert-Bond, 2021). Functional validation of candidate genes in *Ephedra* would provide insight into the conservation, co-option, and evolution of genetic networks. Genetic transformation also complements transcriptomic analyses (Zumajo-Cardona and Ambrose, 2022) by providing an avenue to test the function of candidate genes emerging from gene expression analysis.

While short compared to other gymnosperms, the 2–4-year generation time of *Ephedra* species may pose challenges for applying VIGS in reproductive tissues. Within the scope of our study, photobleaching continued beyond three months, but the full duration of gene silencing remains to be tested. However, there are proven alternatives to address the eventual loss of gene silencing. Wounding and injecting previously infected adult plants can help reestablish silencing, as found in other perennials such as *Aquilegia coerulea* (Gould and Kramer, 2007) and *Thalictrum dioicum* (LaRue et al., 2013), and tubers can be treated in Spring ephemerals, accelerating the gap between treatment and flowering (Di Stilio et al., 2010). Additionally, since various *Ephedra* species are amenable to clonal propagation (O’Dowd and Richardson, 1993), infiltration of reproductive explants prior to rooting would circumvent time to reproduction and provide an opportunity to silence candidate genes in reproductively mature plants. In conclusion, the implementation of VIGS in *E. tweedieana* represents an exciting opportunity to expand functional genetic studies to gymnosperms and, in doing so, bridge the gap between angiosperms and the rest of the seed plants.

## Acknowledgements

Thanks to Dr. Dinesh-Kumar for encouraging this study and sharing plasmids pYL156 and pYL192. Mauricio Bonifacino provided the *E. tweedieana* seeds. This work was funded by the Royalty Research Fund, University of Washington (to VSD), AGG was funded by NSF-GRFP, and JTB by UW Biology Department Top Scholar Award.

