## Supplementary materials for "Enabling the Study of Gene Function in Gymnosperms: VIGS in *Ephedra tweedieana*"

SUPPORTING INFORMATION

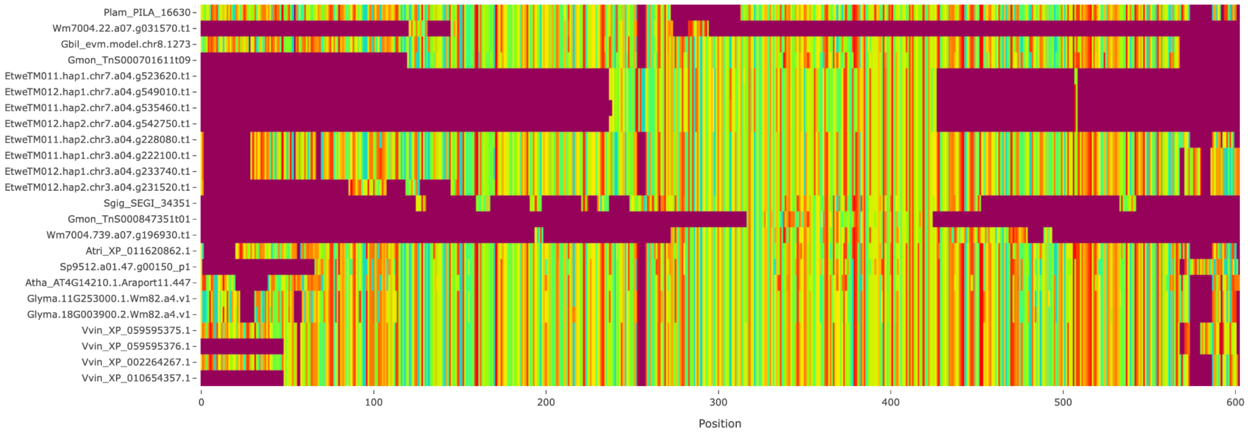


**Figure S1. Multiple sequence alignment (MSA) of the *PHYTOENE DESATURASE* gene family.** The Ephedra Resources web portal ([resources.michael.salk.edu/resources/ephedra_genomes/](resources.michael.salk.edu/resources/ephedra_genomes/)) was searched for the Arabidopsis (AT4G14210) PDS3 ortholog. The two identified orthologs are marked with a bracket, and an arrow points to the one used in the TRV construct.

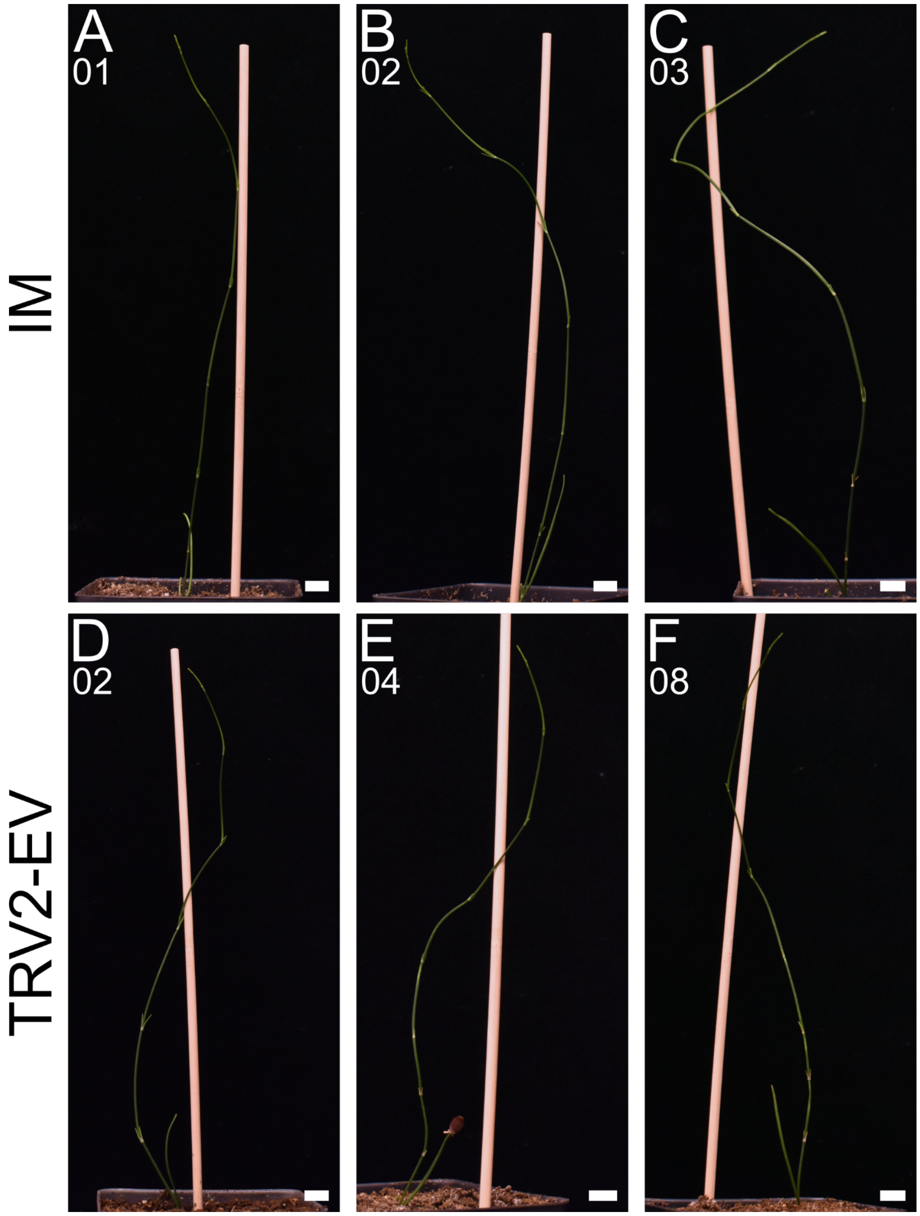


**Figure S2. TRV infection does not affect growth or development.** Each image shows plants three weeks after TRV treatment. a-c) Representative Infiltration-Media (IM) control plants. d-f) Representative TRV*2*-EV control plants displaying similar growth habits as IM control. Scale bar indicates 1 cm.

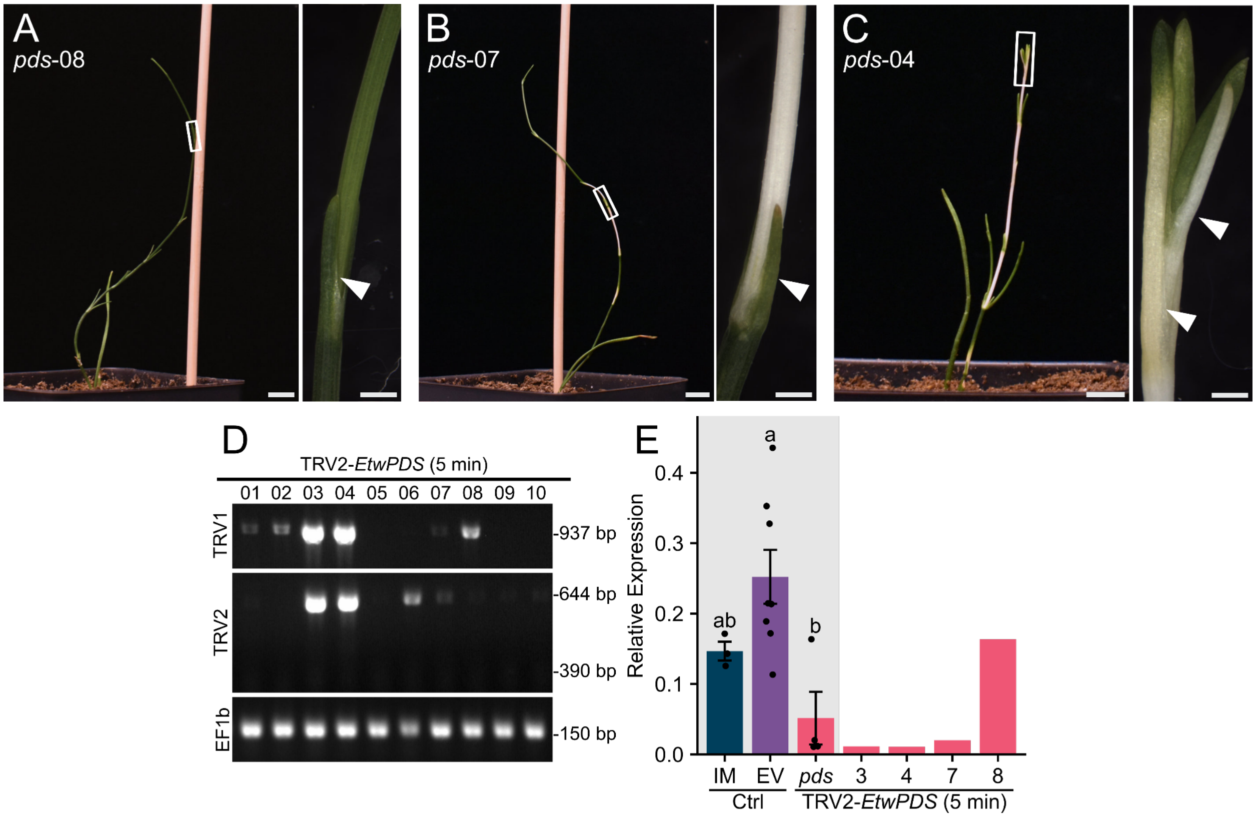


**Figure S3. 5-minute TRV treatment silences *E. tweedieana PDS* and causes tissue bleaching.** Full plant image shows a range of bleaching severity, and the corresponding narrow panel displays magnified detail of the region marked in a white box. a) Empty-vector control plant. b,c) Representative TRV*2-EtwPDS* 5-minute treated plants displaying a range of phenotypes. Scale bars in full plant images indicate 1 cm, while scale bars in magnified panels indicate 1 mm. Arrowheads point to leaves. g) RT-PCR validation of TRV1 RNA and TRV2 RNA presence, including the reference gene *EF1b* as a loading control. h) qPCR validation of *PDS* silencing in bleached tissues of virus-validated plants, relative to EF1b. Different letters indicate statistical significance ($P$<0.05) in a one-way ANOVA ($F_{2,12}$ = 6.672, $P$ = 0.0113) followed by a Tukey HSD post-hoc test (95% family-wise confidence level, $P_{\text{EV vs IM}}$ = 0.241, $P_{\text{EV vs } pds}$ = 0.010, $P_{\text{IM vs } pds}$ = 0.388)

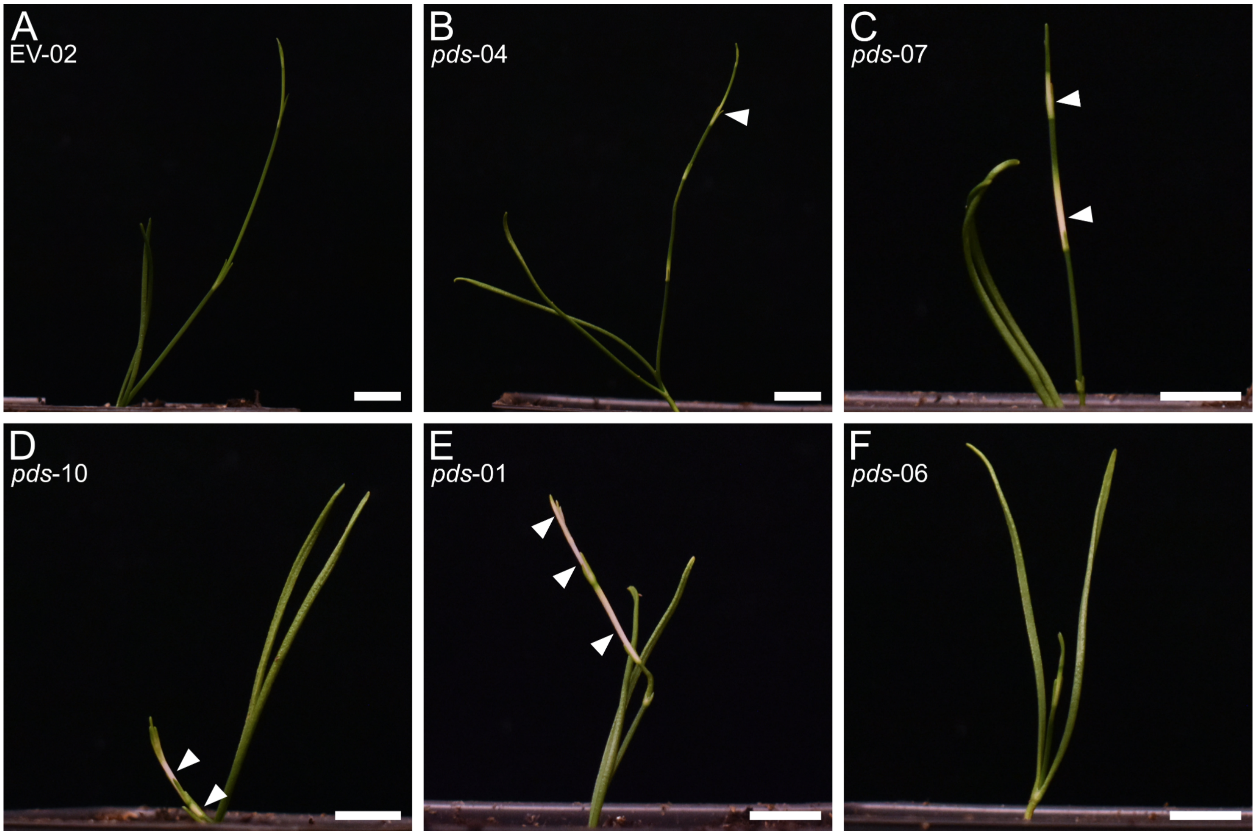


**Figure S4. Photobleaching is observed 1 week after TRV treatment.** Each image shows plants from Figure 2 one week after TRV treatment. a) Empty-vector control plant. b-f) Representative TRV*2-EtwPDS* treated plants displaying a range of phenotypes. Scale bars indicate 1 cm. Arrowheads point to photobleached internodes.

**Table S1. Primers used in this study.**

| Name | Purpose | Sequence |
|---|---|---|
| Etw-PDS-KpnI_F | Amplification and restriction enzyme cloning of EtwPDS | 5'-ttt GGTACC TAGCCGTTTTGACTTTCCAGAG-3' |
| Etw-PDS-XhoI_R | Amplification and restriction enzyme cloning of EtwPDS | 5'-ttt CTCGAG GAGCATTCATTTGAAGCTGCCCA-3' |
| OYL198F | Primer used for RT-PCR of TRV1 | 5'-GTAAAATCATTGATAACAACACAGACAAAC-3' |
| OYL195R | Primer used for RT-PCR of TRV1 | 5'-CTTGAAGAAGAAGACTTTCGAAGTCTC-3' |
| PYL156F | Primer used for RT-PCR of TRV2 | 5'-GGTCAAGGTACGTAGTAGAG-3' |
| PYL156R | Primer used for RT-PCR of TRV2 | 5'CGAGAATGTCAATCTCGTAGG-3' |
| Et-EF1b_F | Primer tested for reference gene qPCR | 5'-GAGGCTAGAGAGGCAACCAA-3' |
| Et-EF1b_R | Primer tested for reference gene qPCR | 5'-AGGCCAGGCATCTCTACACT-3' |
| Et_PDS-qPCR_F_1 | Primer tested for PDS qPCR | 5'-CTGCTGAAGAGTGGATTGGC-3' |
| Et_PDS-qPCR_R_1 | Primer tested for PDS qPCR | 5'-AGCTTTGGTCCCATCTTCTGA-3' |

| | | |
|---|---|---|
| Et_PDS-qPCR_F_2 | Primer tested and used for PDS qPCR | 5’-GACAGTACCCAATTGTGAGCC-3’ |
| Et_PDS-qPCR_R_2 | Primer tested and used for PDS qPCR | 5’-ACAGAACTGCACCCTCCATA-3’ |
| Et_PDS-qPCR_F_3 | Primer tested for PDS qPCR | 5’TCAGAAGATGGGACCAAAGCT-3’ |
| Et_PDS-qPCR_R_3 | Primer tested for PDS qPCR | 5’-AGGCTCACAATTGGGTACTGT-3’ |
